# Elimination of human papillomavirus 16-induced tumors by a mucosal rAd5 therapeutic vaccination in a pre-clinical study

**DOI:** 10.1101/2024.03.11.584519

**Authors:** Molly R. Braun, Jonathan D. Lindbloom, Anne C. Moore, Katherine A. Hodgson, Emery G. Dora, Sean N. Tucker

## Abstract

Therapeutic vaccination can harness the body’s cellular immune system to target and destroy cancerous cells. Several invasive treatments are currently used to eliminate cancerous lesions caused by human papillomaviruses (HPV), however therapeutic vaccination may offer and effective and minimally intrusive alternative. We have developed recombinant, non-replicating human adenovirus type 5 (rAd5) vaccines that encode the HPV16 oncogenic proteins E6 and E7 alongside a molecular dsRNA adjuvant. The potency of these vaccines were examined in a mouse model of HPV tumorigenesis where E6E7-expressing and transformed cells were implanted subcutaneously into C57BL/6 mice. After tumor growth, mice were treated via intranasal administration with E6E7-encoding rAd5 vaccines expressing either a mutant form of E6E7 (rAd5-16/E6E7_m_), or predicted T cell epitopes of E6E7 (rAd5-16/E6E7_epitopes_). Animals receiving therapeutic treatments of rAd5-16/E6E7_m_ and rAd5-16/E6E7_epitopes_ had significant reductions in tumor volume and increased survival compared to animals treated with an empty rAd5 or left untreated. Further, antigen-specific CD8+ T effector memory cells (T_EM_) were observed in the animals treated with E6E7-encoding rAd5, but not in rAd5-empty group. The work described here demonstrates that mucosal rAd5 can be used in a therapeutic capacity to elicit antigen-specific cellular immunity and further identifies a clinical candidate with immense potential for the treatment and prevention of human cervical cancer.

## Introduction

The majority of cervical cancers, as well as oropharyngeal and anogenital cancers, are caused by persistent infection with human papillomavirus (HPV).^1^ There are over 200 types of HPV, with HPV16 and HPV18 being responsible for 71% of cervical cancers.^2,3^ Prophylactic vaccines against the viral L1 capsid protein are currently in use and highly protective against the most common and high risk HPVs (hrHPV), including HPV16 and 18. However, these vaccines are only effective if administered prior to infection and have no therapeutic value if administered once infection has occurred.^4^ HPV is a double-stranded circular DNA virus that largely spreads through sexual contact. Most infections are cleared by the body without intervention, with persistent infection occurring in approximately 10% of patients.^5^ The development of cervical intraepithelial neoplasia (CIN), abnormal changes in the cells lining the cervix, is caused by these persistent infection and is graded 1-3 by level of severity. When left untreated, these abnormal cell pathologies can lead to cervical cancer.^6^ Currently, the standard of care to treat CIN2 and CIN3 involves invasive surgical methods such as ablation or excision.^7^ However, there have been many recent advances in the field of immunotherapy, particularly in therapeutic vaccination, which can provide a non-invasive treatment for cervical cancer. Previous work using therapeutic vaccination to treat cervical cancer has shown efficacy correlated with a robust HPV-specific T cell response.^8^ Therefore, a successfully implemented therapeutic vaccine must be able to stimulate these immune responses.

During infection, viral proteins E6 and E7 modify the cell cycle to promote viral genome amplification using the host cell’s machinery. In some cases, the viral genome may integrate with host genome allowing the continual presence of these proteins.^9,10^ Unchecked expression of E6 and E7 cause unregulated cell entry into S-phase leading to oncogenesis. As E6 and E7 are the causative agents of pre-cancerous and cancerous pathologies, stimulation of cellular immunity specific to these proteins are the target of therapeutic vaccination.^11^ We have developed a mucosal vaccine platform known as Vector-Adjuvant-Antigen Standardized Technology (VAAST), which utilizes recombinant human adenovirus type 5 (rAd5) to express a transgene of interest as well as a molecular dsRNA adjuvant within the same cell.^12-16^ The vaccines manufactured using this platform are formulated into enterically coated tablets that can be administered to humans via swallowing to stimulated a mucosal and systemic immune response.^15^ Our current understanding suggests that after delivery to the ileum^15^, the VAAST rAd5 is taken up by epithelial and resident immune cells. The transgene is expressed and can be displayed on host cells in the context of MHC I or cross-presented on dendritic cells via MHC I and MHC II. Effector T cells recognize the presented vaccine antigens and elicit cell-mediated immunity in the mucosa.^17^ Although rAd5 is delivered to the intestinal ileum, evidence of mucosal crosstalk has been observed in human phase I and phase II clinical trials. In a phase II influenza challenge study, rAd5 encoding the influenza haemagglutinin gene was administered to subjects prior to challenge with influenza virus. The vaccine elicited HA-specific antibodies, mucosal homing B cells, and protected subjects from challenge with influenza.^13^ Additionally, phase I and phase II clinical trials have shown antigen-specific antibodies in saliva and nasal secretions^18,19^ as well as mucosal homing α4β7+ B and T cells.^12,13,15^ Therefore, although this oral tablet vaccine is administered via the intestinal mucosa, there is direct evidence of mucosal crosstalk.

HPV16 does not infect animals nor lead to the mucosal tumorigenesis as seen in humans.^20^ There are new methods in development that model orthotopic HPV tumors; however, these studies are technically challenging and not widely available.^2^ Here we describe the use of rAd5, engineered to encode HPV16 E6 and E7 in a therapeutic capacity. In a proof-of-concept experiment, we employed a commonly available HPV tumorigenesis model that uses the TC-1 cell line, a murine C57BL/6 lung cell line generated by transduction with HPV16 E6 and E7 as well as variant H-ras activated by the G12V mutation.^21,22^ These cells can be cultured *in vitro* and implanted subcutaneously into C57BL/6 mice, providing a surrogate model to test therapeutic vaccinations.^21,23^

In this study, we examined the antitumor effects of mucosal therapeutic vaccination with rAd5 encoding the oncogenic HPV16 genes E6 and E7. We found that the VAAST HPV16-specific rAd5, when administered to tumor-bearing mice, led to a significant reduction in tumor volume and increased survival. Relative percentages of CD4 and CD8 T cells remained consistent between the groups treated with E6/E7 antigens compared to a group treated with an empty vector. No significant differences were observed between naïve, central memory, or effector memory T cells percentages. However there was a significant increase in E7-specific CD8+ T cells, which was largely made up by T effector memory cells. Our results indicate that these mucosal E6/E7 expressing rAd5 vectors are efficacious and immunogenic in a mouse model of HPV-derived tumorigenesis and may represent an effective, non-invasive, and potent therapeutic for the treatment of HPV-related cancers.

## Methods

### rAd5 generation

The transgenes expressed by the adenovirus type 5 (rAd5) vaccines were generated based on the published sequence of HPV16 E6 (GenBank Accession Number ANY26540.1) and E7 (GenBank Accession Number AIQ82815.1). For rAd5-16/E6E7_m_, E6 mutations L57G, E154A, T156A, Q157A, and L158A and E7 mutations H2P, C24G, E46A, and L67R were introduced. A furin cleavage site separates the E6 and E7. For rAd5-16/E6E7_epitopes_, amino acids 18-26, 42-56, 126-143 of E6 and 4-23, 62-77, and 81-90 of E7 were included without spacers. The transgene sequences described above were inserted into the E1 region of a recombinant plasmid containing the rAd5 genome lacking the E1 and E3 genes. A third rAd5 construct was used that did not contain a transgene sequence (rAd5-empty). A sequence encoding the molecular dsRNA adjuvant was included downstream of the transgene region. Both E6/E7 and the dsRNA are under control of CMV promoters. The rAd5 vaccines were generated and propagated as previously described.^24^

### TC-1 cell line and tumorigenesis

TC-1 cells (ATCC, USA) were grown as a monolayer at 37°C with 5% CO_2_ in RPMI 1640 supplemented with 2 mM L-glutamine (31870-025, Thermofisher), 1 mM sodium pyruvate (11360-039, Thermofisher), 0.1 mM non-essential amino-acids (11140-035,Thermofisher), 50 μM β-mercaptoethanol (31350-010, Thermofisher), 1% penicillin/streptomycin (15140-122, Thermofisher) and 10% fetal bovine serum (P30-3306, Pan Biotech). For tumor implantation, cells were detached from flask with trypsin-0.05% EDTA (25300054, Thermofisher) and 1×10^6^ TC-1 cells in 200 μL of RPMI 1640 without phenol red were infected into the right flank of 50 C57BL/6JRj mice (female, 7 weeks old at reception, Janiver Labs). Animals were randomized by mean tumor volume when tumors reached 20-60 mm^3^. 40 of the mice were divided into four groups of 10 animals using Vivo Manager software (Biosystemes, France). Animals were monitored for health parameters and tumor volume. All work described was performed by Oncodesign Services (Dijon, France).

### Immunization

Theraputic vaccine treatments were started on the day of randomization and repeated twice more, seven days apart (Q7Dx3). Groups received 1×10^8^ IU/animal of rAd5_empty_, rAd5-16/E6E7_m_, rAd5-16/E6E7_epitopes_, or were untreated via intranasal administration. All work described was performed by Oncodesign Services (Dijon, France).

### Flow cytometry

Blood was collected by jugular vein puncture on days 14 and 21. Prior to staining, red blood cells (RBCs) were lysed with Versalyse lysing buffer (A09777, Beckman coulter) for 10-15 minutes. Cells were stained with viability dye Viakrom 808 (Beckman Coulter), followed by Staining with Dextramer E7_49-57_ APC (Immudex, Allele H-2Db). Cells were then stained with CD45 BV605 (103155, Biolegend), CD3 PE (130-120-160, Miltenyi), CD4 (130-123-894, Miltenyi), CD8 BV785 (100750, Biolegend), CD62L (BV421 104436, Biolegend), and CD44 VioBright FITC (130-120-213, Miltenyi). Cells were fixed with Cytofix buffer (554655, BD Biosciences) then run on a flow cytometer. All work described was performed by Oncodesign Services (Dijon, France).

## Results

### Rational Design of Therapeutic Vaccines for HPV16

The E6 and E7 proteins of HPV16, when unchecked, can cause dysregulation of the cell cycle and tumorigenesis. The molecular virology of rAd5 delivery allows only a single round of target cell transduction. Nevertheless, it is inadvisable to deliver oncogenic genes during vaccination. Therefore, two safety approaches were taken. In the first approach, an rAd5 vaccine, rAd5-16/E6E7_m_, was generated with mutations that disrupt E6’s ability to bind p53 and PTPN13 and E7’s ability to bind Rb and mi2β.^25^ In a second approach, a construct, rAd5-16/E6E7_epitopes_, was generated to express the minimal epitopes required for immunogenicity and efficacy, utilizing the Immune Epitope Database & Tools, a resource that uses MHC class I binding, peptide processing, and immunogenicity predictions to identify consequential epitopes.^26^ As a comparator, a third vaccine which only contains the molecular dsRNA adjuvant, rAd5-empty, was used in this study (Figure 1A).

**Figure 1:**
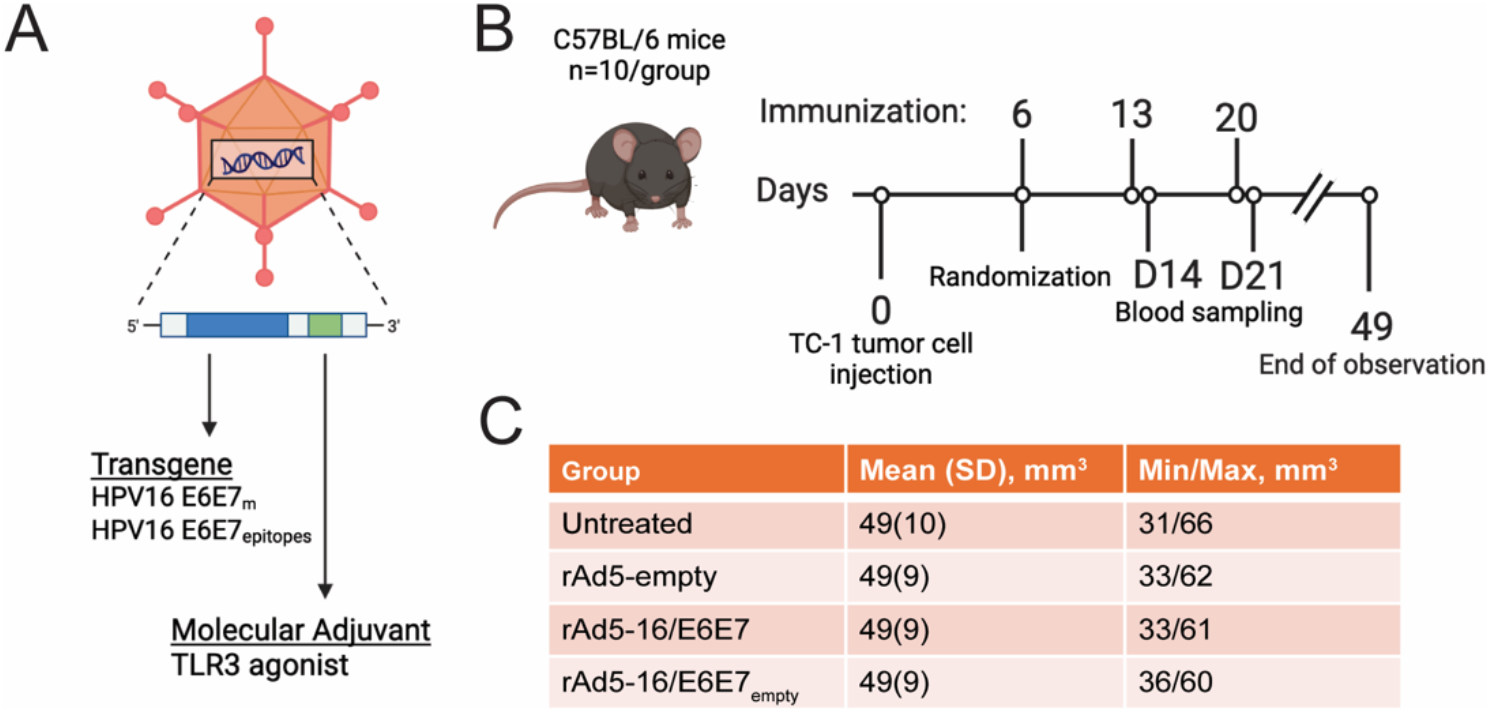
Study design for immunogenicity and efficacy of rAd5 vaccines in TC-1 tumor model. A) Illustration of the rAd5 vector used during vaccination. The transgene region (blue) represents the antigen included in the construct upstream of the molecular dsRNA adjuvant (green). (B) Schedule of tumor initiation, vaccination, blood sampling, and tumor observation time. (C) Tumor size at the day of randomization (study day 6)

### Mucosal application of antigen-specific rAd5 reduces tumor size

On study day 0, TC-1 tumors were induced by subcutaneous injection of 1×10^6^ TC-1 cells into the right flank of C57BL/6 mice. Tumors grew until a mean volume of 20-60 mm^3^ was reached. Animals were randomized on day 6 into groups with an average tumor volume of 49 mm^3^ (Figure 1B, C). Mice were then treated with rAd5-empty, rAd5-16/E6E7_m_, or rAd5-16/E6E7_epitopes_ (Figure 1B,C) by intranasal application three times, seven days apart, or left untreated.

After randomization and initial treatment on day 6, tumors continued to grow in size until day 13, at which point there was no significant difference in tumor size between treatment groups (Figure 2A,B, supplemental figure 1). Animals received additional treatments on day 13 and day 20 (Figure 1B). After day 13, tumors in the untreated group and the group treated with rAd5-empty continued to increase in volume until ethical standards were met for euthanasia (Figure 2C) whereas tumor volume in the groups treated with rAd5-16/E6E7_m_ or rAd5-16/E6E7_epitopes_ began to decrease in size through the end of the study at day 49 (Figure 2A, Supplemental Figure 1). At day 29, the last day that 80% of animals were present in each group, the rAd5-16/E6E7_m_ and rAd5-16/E6E7_epitopes_ groups had significantly smaller tumors with the group treated with rAd5-16/E6E7_m_ having nearly undetectable tumors at this timepoint (Figure 2B). The effect of rAd5-16/E6E7_m_ was particularly pronounced as tumor completely regressed in all animals with lastly regression in 5/10 animals (Supplementary Figure 1).

**Figure 2:**
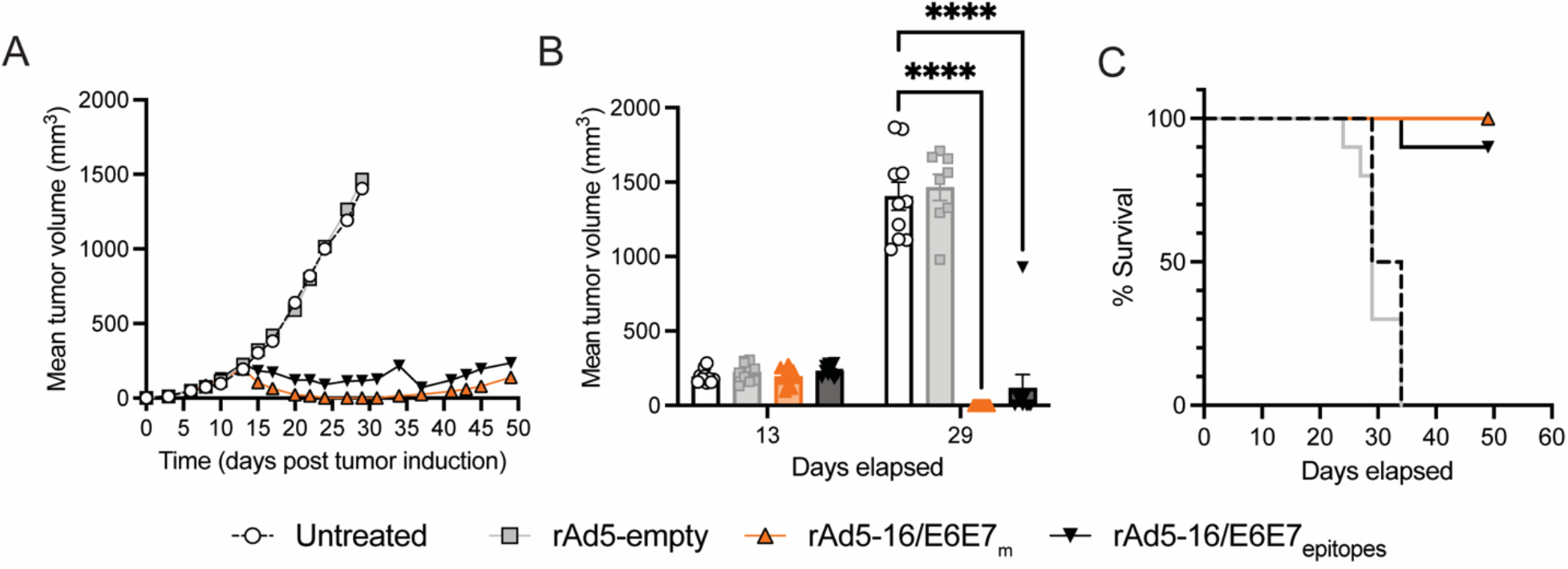
E6-E7 specific rAd5 vaccines prevent tumor growth in mice bearing subcutaneous TC-1 tumors. (A) Mean tumor volume per group through study. Curves are shown for data points were at least 80% of animals were present. (B) Mean and SEM of tumor volume at indicated days. n=10 mice per group, 2-way ANOVA (C) Survival curve indicating time until ethical criteria for euthanasia was met.

### Mucosal application of antigen-specific rAd5 generates antigen-specific T_EM_ cells

As oncogenesis is driven by E6 and E7, which are intracellular targets, cellular immunity was characterized by flow cytometry one day after the second and third treatment, on days 14 and 21, to look for changes in T cell distributions as well as generation of antigen-specific T cells. Blood was sampled from animals from the groups treated with rAd5, but not the untreated group. On day 14, the percentage of CD45+ cells was highly variable between samples; however, despite this variability, the percentage of CD3+ cells within the CD45+ population was consistent between groups with the exception of a statistically significant increase in overall percentage of T cells in the group treated with rAd5-16/E6E7_m_ compared to rAd5-empty at day 14 (Figure 3A). This difference was not present at day 21, one day after second treatment. Among CD3+ T cells, there were no differences in overall distribution of either CD8+ T cells or CD4+ T cells at either sampling day between the various groups (Figures 3B,C).

**Figure 3:**
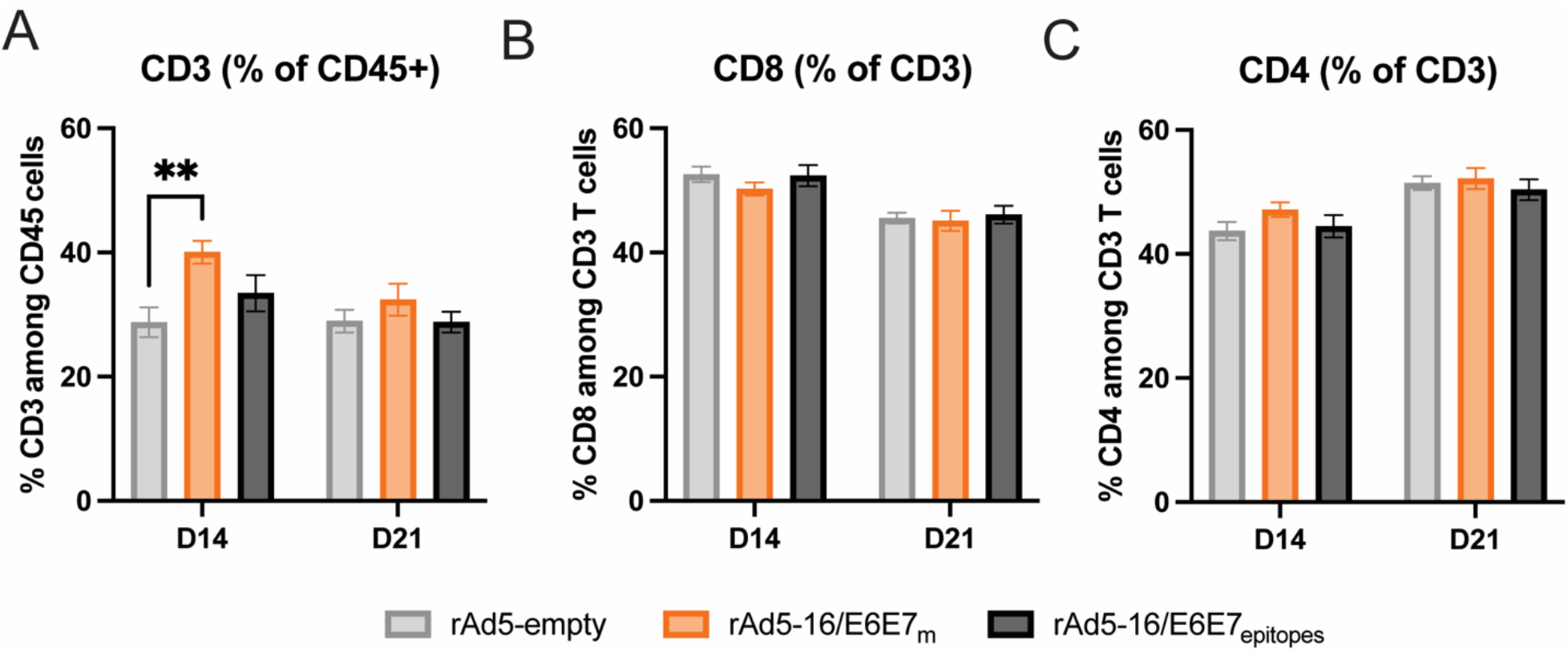
T cell distributions remain similar between animals treated with rAd5 vaccines. T cells in the blood of animals sampled one day after D13 and D20 vaccinations. (A) Total T cells as a percent of lymphocytes (CD45+). (B) CD8+ T cells as a percent of CD3+. (C) CD4+ T cells as a percent of CD4+, were measured by flow cytometry. n=10 mice/group, mean and SEM, 2-way ANOVA.

To further characterize the cellular immune response, we next examined the cellular markers CD44 and CD62L in the T cell population to distinguish between naïve, central memory (T_CM_), and effector memory (T_EM_) T cells. There were no significant differences or consistent trends in distribution between CD8+ T cell subsets (Figure 4A-C) or CD4+ T cell subsets (Supplementary Figure 2). As T_EM_ cells are the main drivers of cellular cytotoxicity towards cancerous cells, we sought to understand if E7-specific T cells were being produced by rAd5 treatments using a Dextramer with a known E7_49-57_ H-2 Db allele (DexE7) bound to MHC class I. On day 14, after two treatments, the rAd5-16/E6E7_m_ group had a significant increase in E7 specific T cells compared to the rAd5-empty group. At day 21, after three treatments, both the rAd5-16/E6E7_m_ group and the rAd5-16/E6E7_epitopes_ group had significantly increased percentages of DexE7 specific CD8+ T cells with rAd5-16/E6E7_m_ having a significantly higher response to both rAd5-empty and rAd5-16/E6E7_epitopes_ (Figure 4D). Of the DexE7+ CD8+ T cells, most cells were T_EM_ cells, representing a critical cell population for cell-mediated control of HPV tumorigenesis (Figure 4E).

**Figure 4:**
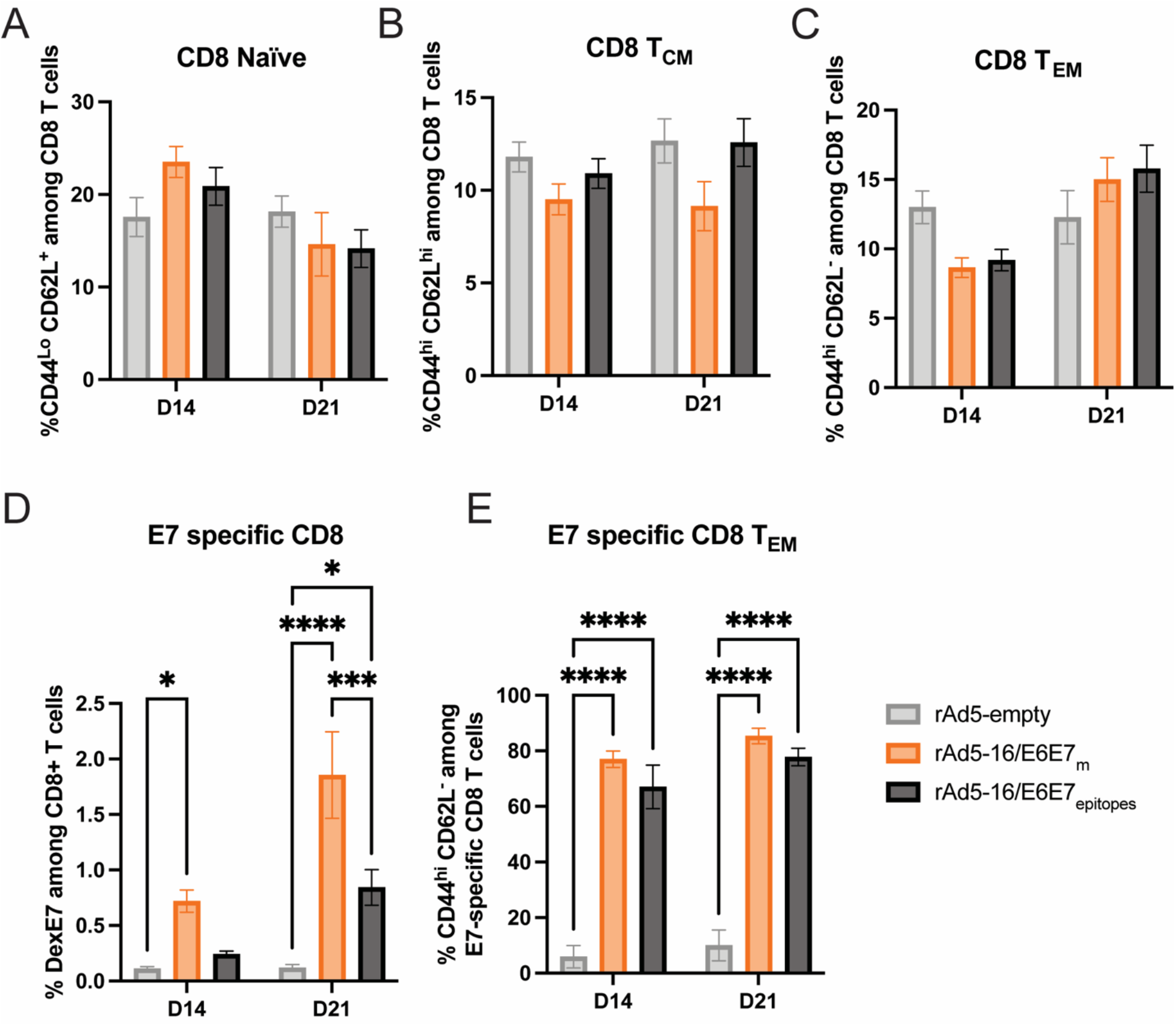
T_EM_ make up most antigen-specific T cells. (A) Naïve, (B) central memory, or (C) effector memory CD8 T cells did not alter in relative percentage as a result of transgene expressed. Small differences were not statically significant n=10 mice/group, mean and SEM, 2-way ANOVA. (D) Percentage of CD8+ T cells specific to HPV16 E7. (E) Percentage of E7-specfic T cells among E7 specific cells. n=10 mice/group, mean and SEM, 2-way ANOVA.

## Discussion

Therapeutic vaccination, designed to stimulate the body’s cell-mediated immune response to cells transformed by HPV infection, may harness the host’s immune system to create a potent cell-mediated response and may also provide a less invasive, more widely available treatment for CIN2/3. A thermostable oral tablet vaccine that can be used therapeutically to activate these necessary immune responses would be greatly impactful in the treatment and prevention of cervical cancer. As HPV screening systems are put in place, infections of hrHPV can be caught earlier. Non-invasive treatments, like the one examined here, may even used at the point of infection identification prior to diagnosis with CIN2/3 to prevent the progression to from CIN1 to cervical cancer.^27^

In this report, we evaluated a novel HPV16 E6/E7 vaccine that contains previously described mutations that inhibit E6 and E7 oncogenic properties or contains only the predicted immunodominant epitopes to increase safety.^25^ We demonstrate that mucosal vaccination via an intranasal route in mice was able to reduce tumor burden in a subcutaneous model of HPV tumorigenesis. All animals receiving E6E7-containing rAd5 vaccines exhibited tumor control, and in some cases, complete regression of the tumor, through the end of the study on day 49. This effect was especially pronounced in the rAd5-16/E6E7_m_ group with complete regression of tumors in all mice and lasting regression in half of the mice. In contrast, animals in the untreated or rAd5-empty group met ethical criteria for euthanasia by day 34.

The generation of antigen-specific cytotoxic CD8 T cells plays a critical role in immunotherapies and thus, it is critically important that any therapeutic vaccine be able to generate this type of response.^28^ We sought to understand how T cell dynamics changed in response to antigen-specific vaccination. Administration of the VAAST rAd5 did not appear to alter the overall distribution of CD4 and CD8 T cells, but rAd5 encoding E6E7 genes were able to generate antigen specific CD8 T_EM_ cells in the periphery, with rAd5-16/E6E7_m_ generating a more pronounced response than rAd5-16/E6E7_epitopes_. In this study, we did not examine mucosal homing T-cells, however our group has previously shown the generation of mucosal homing α4β7+ T cells after vaccination in human clinical trials.^12-14^ Future studies will investigate the generation of these cells as well as the presence of tumor-infiltrating lymphocytes, another key indicator of tumor regression.

Several therapeutic vaccines for HPV have completed placebo-controlled clinical trials and have largely been administered via injection or electroporation. A review of different vaccine modalities using variations of HPV16 and HPV18 E6 and E7 as antigens, observed a total overall proportion of regression from CIN2/3 to CIN1 at 0.54 compared a placebo group at 0.27.^27^ These numbers represent a meaningful reduction in advancement to cervical cancer, yet room for improvement remains. Studies with an electroporated DNA vaccine candidate suggested that the ability to elicit potent antigen-specific T cell responses correlated with histopathological regression and a reduction in HPV viral DNA detection.^29^ A study monitoring women with high grade squamous intraepithelial lesions (HSIL), a term used to characterize CIN2 and CIN3, indicated that α4β7+ CD8 T-cells are better able to enter the cervical mucosa and are associated with cervical lesion clearance compared to those excluded from the epithelium.^30^ In this report, we demonstrate the generation of antigen-specific CD8 T cells induced by therapeutic vaccination via a mucosal vaccine while also demonstrating a reduction in tumor volume. Thus, this the VAAST rAd5 platform provides a promising technology to create a therapeutic vaccine to treat HPV derived cervical dysplasia.

---

**Supplemental Figure 1:**
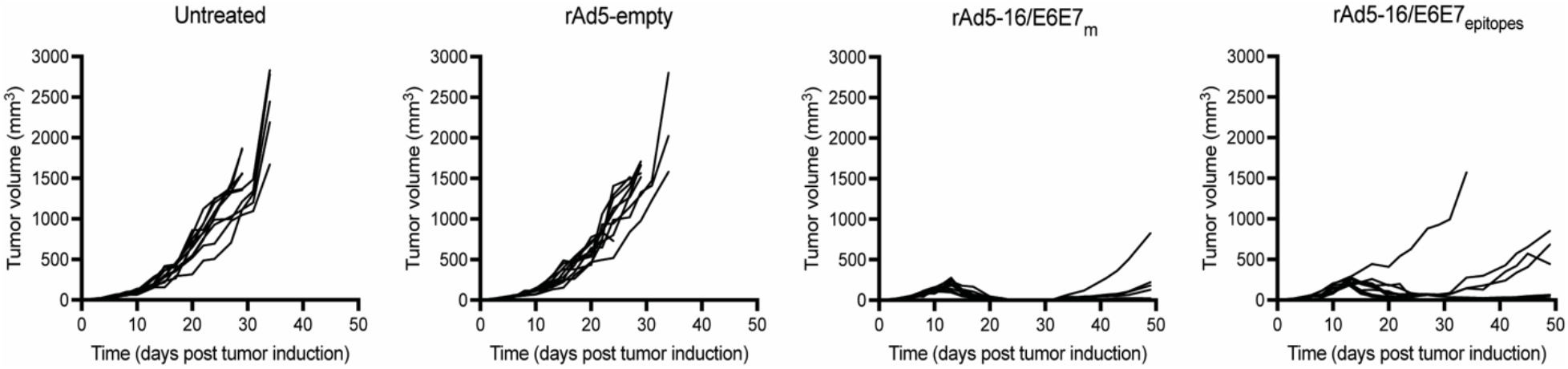
Individual tumor volumes of C57BL/6 mice bearing subcutaneous TC-1 tumors.

**Supplemental Figure 2:**
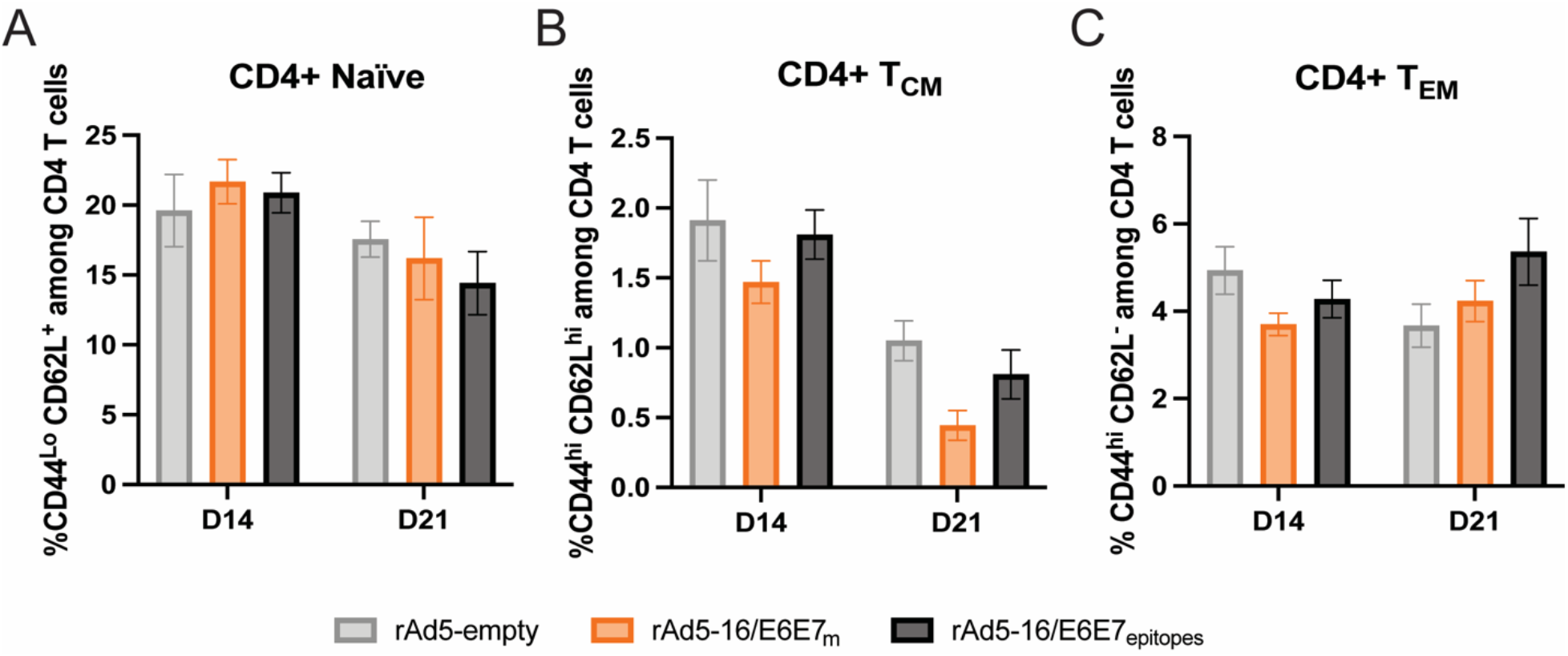
CD4+ T cell distributions remain similar between animals treated with rAd5 vaccines. A) Naïve, (B) central memory, or (C) effector memory CD4+ T cells did not alter in relative percentage as a result of transgene expressed. Small differences were not statically significant n=10 mice/group, mean and SEM, 2-way ANOVA.

